# Regulation of Cell-Type-Specific Transcriptomes by miRNA Networks During Human Brain Development

**DOI:** 10.1101/412510

**Authors:** Tomasz J Nowakowski, Neha Rani, Mahdi Golkaram, Hongjun R Zhou, Beatriz Alvarado, Kylie Huch, Jay A West, Anne Leyrat, Alex A Pollen, Arnold R Kriegstein, Linda R Petzold, Kenneth S. Kosik

**Affiliations:** Eli and Edythe Broad Center of Regeneration Medicine and Stem Cell Research, University of California, San Francisco; Department of Anatomy, University of California, San Francisco; Department of Psychiatry, University of California, San Francisco; Neuroscience Research Institute University of California, Santa Barbara; Department of Molecular, Cellular, and Developmental Biology, University of California, Santa Barbara; Department of Biological Sciences & Bioengineering, Indian Institute of Technology, Kanpur, 208016, India; Department of Mechanical Engineering, University of California, Santa Barbara; Department of Neurology, University of California, San Francisco; New Technologies, Fluidigm Corporation, South San Francisco; Department of Computer Science, University of California, Santa Barbara

## Abstract

MicroRNAs (miRNAs) regulate many cellular events by regulating hundreds of mRNA transcripts. However, it is unclear how miRNA-mRNA interactions are contextualized into the framework of transcriptional heterogeneity among closely related cells of the developing human brain. By combining the multiple complementary approaches, AGO2-HITS-CLIP, single-cell profiling and bipartite network analysis, we show that <u>the</u> miRNA-mRNA network operates as functional modules related to cell-type identities and undergo dynamic transitions during brain development.

---

The interactions between miRNAs and mRNAs play essential roles during mammalian development. Individual miRNAs have been functionally implicated in a variety of functions related to specific tissues, anatomical and cellular compartments, evolutionary relationships, and developmental time points^1-4^. In the developing mouse cortex, ablation of *Dicer1* further highlighted key roles for miRNAs in cell fate transitions^5,6^. However, systematic analysis of the expression specificity of miRNAs and their interaction with mRNAs remains challenging. Bioinformatic predictions based on miRNA seed regions are overly broad, requiring direct empirical studies to reveal the specific interactions between miRNAs and mRNAs that are essential for normal development. In addition, systematic analysis of miRNAs at the single cell level may add richness to classifications of cell identity and inference of function by representing a key post-transcriptional modality^7,8^. By leveraging insights from three complementary approaches: RNA isolated by crosslinking immunoprecipitation (HITS-CLIP), single cell quantitative RT-PCR (sc-qPCR), and single-cell RNA sequencing (scRNA-seq), we sought to characterize the miRNA-mRNA interactions during *in vivo* human brain development and to contextualize these networks in the framework of developmental transitions and cell identity. These technologies uncovered dynamic networks involving cell type specific enrichment of individual miRNA expression across diverse cell types as well as miRNA target acquisition and loss in which the population of targeted mRNAs keeps pace with the dynamics of tissue development, cell diversity, and lineage progression through human brain development.

Expression profiling of miRNAs using bulk tissue samples suggested developmental regulation of miRNA expression^9^; however, the extent to which these profiles change during developmental cell fate transitions has not been comprehensively addressed at single cell level except among distantly related cell types^7^. We developed a protocol for combined miRNA and mRNA sc-qPCR in single cells (Fig. 1a and Supplementary tables 1-2) which leverages a microfluidic platform to perform automated cell capture, reverse transcription and targeted preamplification of mRNA and miRNA. Target mRNAs were selected based on the specificity of their expression across diverse cell types of the developing human brain (Extended Data Fig 1a) as determined in Pollen AA et al.,^10^.

**Fig. 1.**
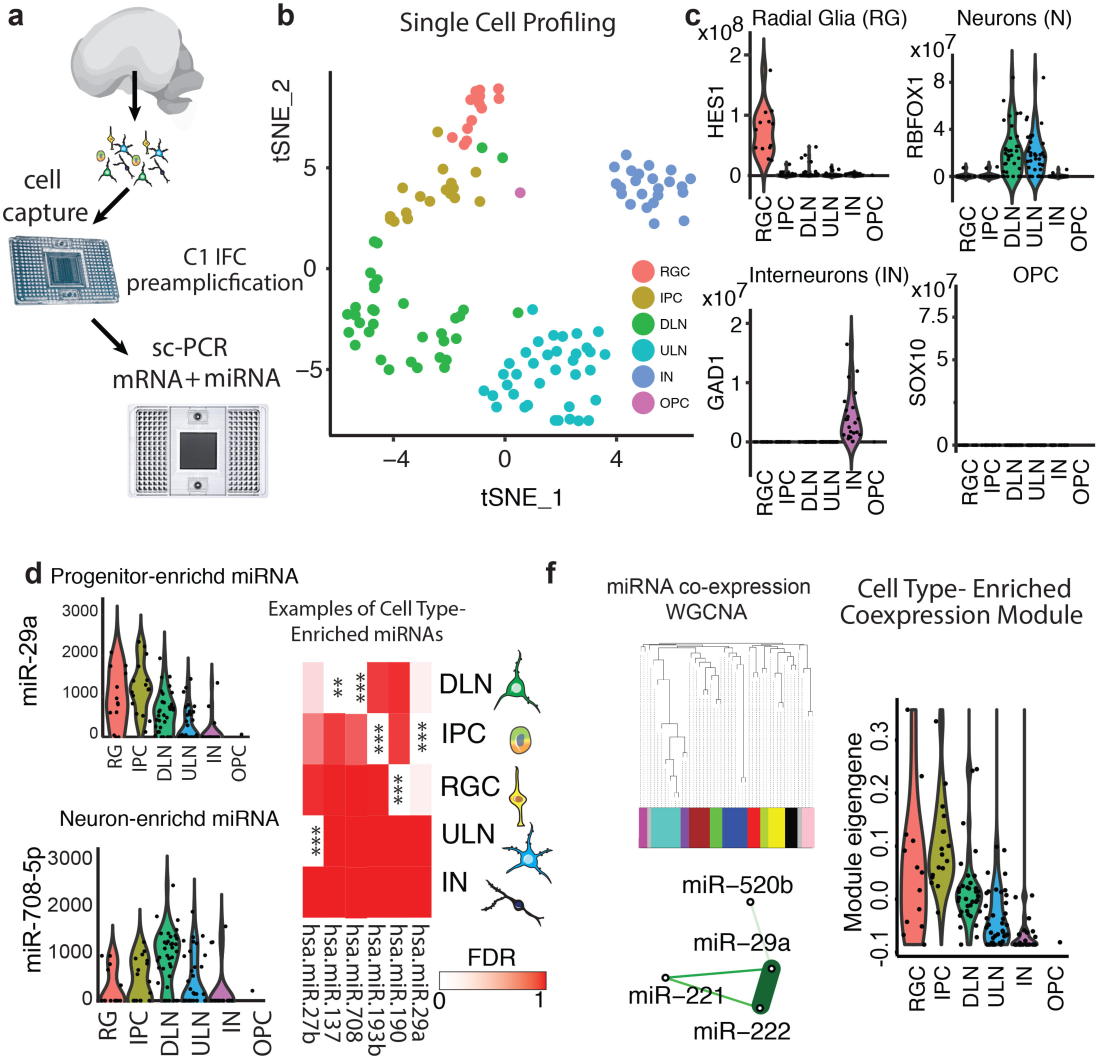
Single-cell miRNA Expression Profiling Reveals Patterns of Cell-type Enrichment. **(a-c)** sc-qPCR profiling of mRNA and miRNA abundance in the same cell (a). **(b)** tSNE plot of single-cell data generated. RGC– radial glia cells, IPC - intermediate progenitors, ULN-upper layer neurons, DLN-deep layer neurons, IN– interneurons, OPC-oligodendrocyte progenitor cell. **(c)** Marker gene expression. **(d)** Violin plots show miRNAs with cell-type enrichment. Heatmap represents examples cell-type enriched miRNAs profiled. ^**^FDR<0.01, ^***^FDR<0.001, one-tailed U-test. **(f)** Weighted gene co-expression network analysis reveals modules of co-expressed miRNAs across single-cells profiled, including the example module enriched in progenitor cells.

We used this approach to profile single cells isolated from primary human cortical tissue samples at gestational week (GW) 15 and GW17. Using cell type specific marker expression, we inferred the identities of the cells as radial glia, intermediate progenitors, upper and deep cortical layer neurons, interneurons (Fig. 1a-c, Supplementary Fig. 1), or oligodendrocyte precursors. In addition, by projecting miRNA abundance across the same cells, we calculated an expression enrichment score for every miRNA (Fig. 1d) and found striking enrichment of multiple miRNAs in distinct cell types, including the early born neurons captured from the germinal zone, suggesting that dynamic changes in miRNA abundance occur simultaneously with changes in mRNA abundance during neuronal differentiation (Fig. 1E, fig. S1). Moreover, grouping individual miRNAs based on correlated expression patterns further revealed that some highly-specific miRNA co-expression modules operate within specific cell types, whereas others are broadly distributed across multiple cell-types (Fig. 1e-f, Supplementary Fig. 1). These results demonstrate precise regulation of miRNAs, even among closely related developmental cell populations during cell fate transition in the developing human cortex.

To identify the genes engaged in miRNA-mRNA interactions during human brain development *in vivo*, we performed high-throughput sequencing of RNA isolated by crosslinking immunoprecipitation (HITS-CLIP)^11^ with AGO2 in primary developing human brain tissues (Fig. 2, Supplementary Fig. 2, Supplementary Table 2-5). AGO2 HITS-CLIP provides the strongest confidence for target identification. Gene enrichment analysis highlighted regulation of transcription, chromatin modification, and signaling pathways among miRNA targets (Supplementary Table 6). Surprisingly, many target genes are well-established marker genes^10,12,13^ (Fig. 2c, Supplementary Fig. 3) and important regulators of neurogenesis, migration, axonogenesis, synaptogenesis, and neuronal subtype specification (Fig 2b), suggesting that miRNAs regulate key developmental decision during excitatory lineage progression by targeting key cell type specific genes (Fig. 2c, Supplementary Table 7).

**Fig. 2:**
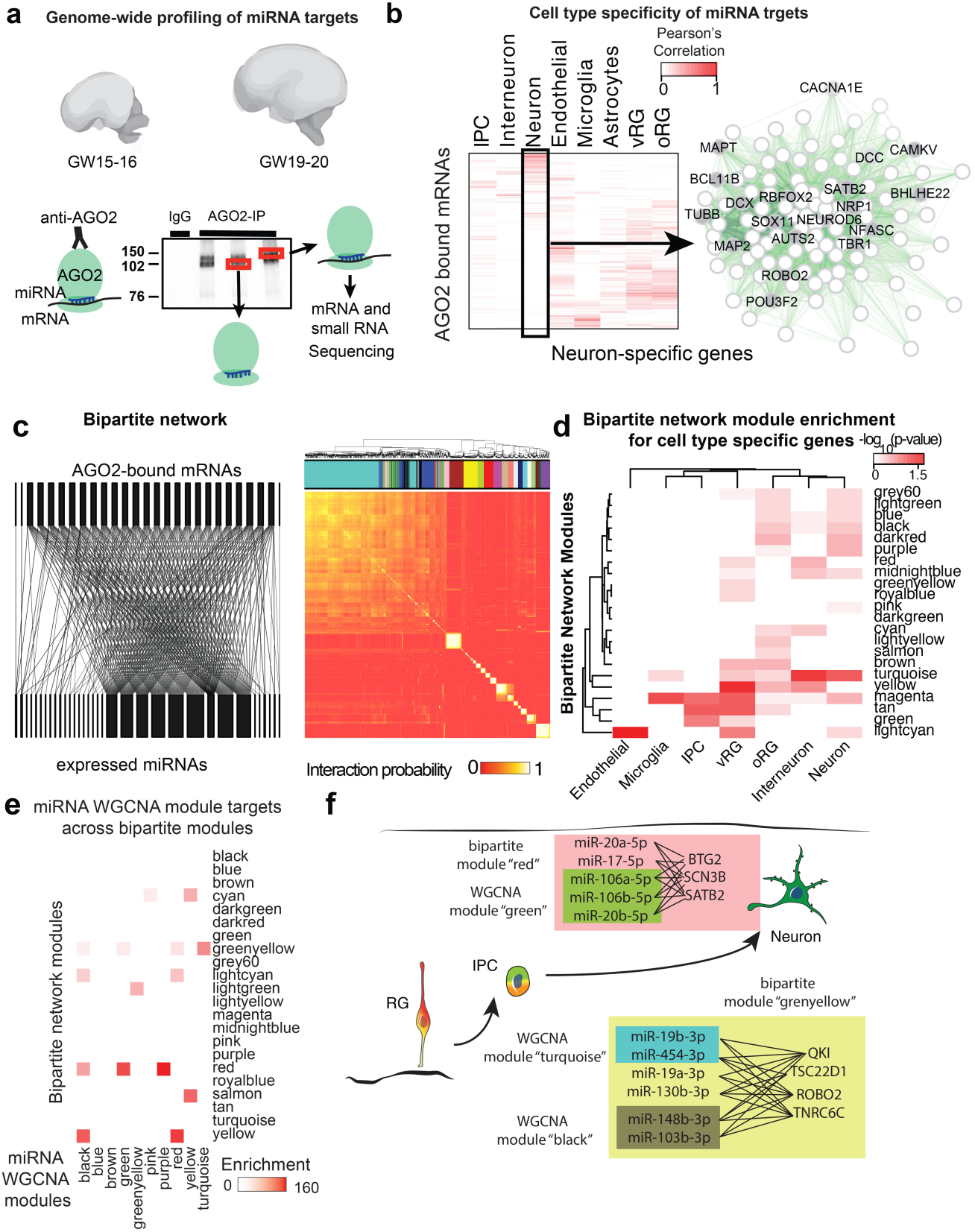
High Throughput Profiling of miRNA-mRNA Interactions. **(a)** Schematic of HITS-CLIP analysis. **(b)** Cellular specificity of target genes^10,12^. Network plot shows neuron-specific genes targeted by miRNAs. **(c)** Bipartite network analysis of miRNA-mRNA interactions. **(d)** Enrichment of bipartite modules for cell-type specific transcripts. **(e)** Enrichment of miRNA coexpression networks across bipartite network modules. Enrichment score represents –log_10_(p-value). **(f)** Examples of interactions between coexpressed miRNAs and their targets within bipartite network modules.

Interactions between miRNAs and their direct target mRNAs can be presented as a bipartite network (Fig. 2c) because the principle regulatory interactions are between miRNAs and mRNAs rather than regulating themselves. As expected, single miRNAs target many mRNAs and single mRNAs are targeted by far fewer miRNAs (Supplementary Fig. 4a-b). A bipartite community detection algorithm^14^ revealed modules of miRNA-mRNA interactions with significantly greater intra-modular than inter-modular connectivity (Fig. 2c, Supplementary Fig. 5-6, Supplementary Table 8, see Methods for details). Interestingly, modules tend to recruit several low-abundant miRNAs or few high-abundant miRNAs (Supplementary Fig. 4, 7), suggesting that the implementation of a miRNA mediated function targets either with one or a very few abundant miRNAs or with multiple low abundance miRNAs. To contextualize these interactions in the framework of cell diversity we projected cell type specificity information, calculated from published scRNA-seq datasets, against bipartite graph modules (Fig. 2d). Our analysis revealed striking enrichment for cell-type-specific transcripts among bipartite graph modules, suggesting that miRNAs acquire targets according to the cognate transcriptional landscape of individual cell-types.

Recent studies suggest perturbations in miRNA expression may contribute human developmental neuropsychiatric disorders^15,16^, but the specific molecular consequences remain poorly understood. Interestingly, we found that miR-137 and miR-218, which have recently been implicated in human disease, showed enrichment in a subset of deep layer maturing pyramidal neurons (Supplementary Fig. 1b-c and Supplementary Fig. 6). In addition, genes implicated in autism spectrum disorders (ASD)^17^, are enriched in the greenyellow bipartite module, which includes miR137 and miR-218 targets (Supplementary Fig. 6). Moreover, genes targeted by miR-137 in developing brain are vastly different than targets identified in adult human brain tissue^18^, suggesting dynamic regulation of the miR-137 target landscape (Supplementary Table 9). Understanding cell-type-specific miRNA expression profiles and their respective targets may highlight cellular patterns of selective vulnerability to disorders affecting miRNA expression by highlighting gene regulatory networks that might be perturbed in disease states.

To identify miRNA-mRNA interactions networks involved in human cortical neurogenesis *in vivo*, we mapped miRNA co-expression networks from sc-qPCR (Fig. 1e-f, Supplementary Fig. 1b-c) onto bipartite co-regulatory modules inferred from HITS-CLIP (Fig. 2d). This analysis revealed a striking enrichment of targets of co-expressed miRNAs among bipartite network modules. For example, targets of miRNA co-expression module ‘brown’ (Supplementary Fig. 1), which is enriched in IPCs and neurons, are enriched in bipartite module ‘yellow’, which is strongly enriched for radial glia specific genes (Fig. 2e-f). Our analysis provides an unbiased view of miRNA regulatory networks involved in human cortical neurogenesis.

Next, we compared the abundance of miRNAs at two stages of development - GW15-16.5 and GW19-20.5 and found 69 differentially expressed miRNAs between these two stages including recently evolved miRNAs (Fig. 3a-b, Supplementary Figs. 8-10, and Supplementary Table 10-11). We also identified miRNAs enriched in the occipital lobe compared to the frontal lobe (Supplementary Fig. 11-13, Supplementary Table 4), suggesting, along with recent studies^19^, that miRNAs may regulate regionally divergent transcriptional states in the developing human cortex. By independently performing bipartite network analysis for samples at these two stags, we found a striking preservation of most co-regulatory modules, and a set of distinct interactions present predominantly at one stage (Fig. 3c, Supplementary Fig. 14, Supplementary Tables 8 and 12). Together, our findings suggest that miRNA-mediated regulation forms a developmentally dynamic network of interactions related to cell type, developmental stage, and cortical area specificity.

**Fig. 3.**
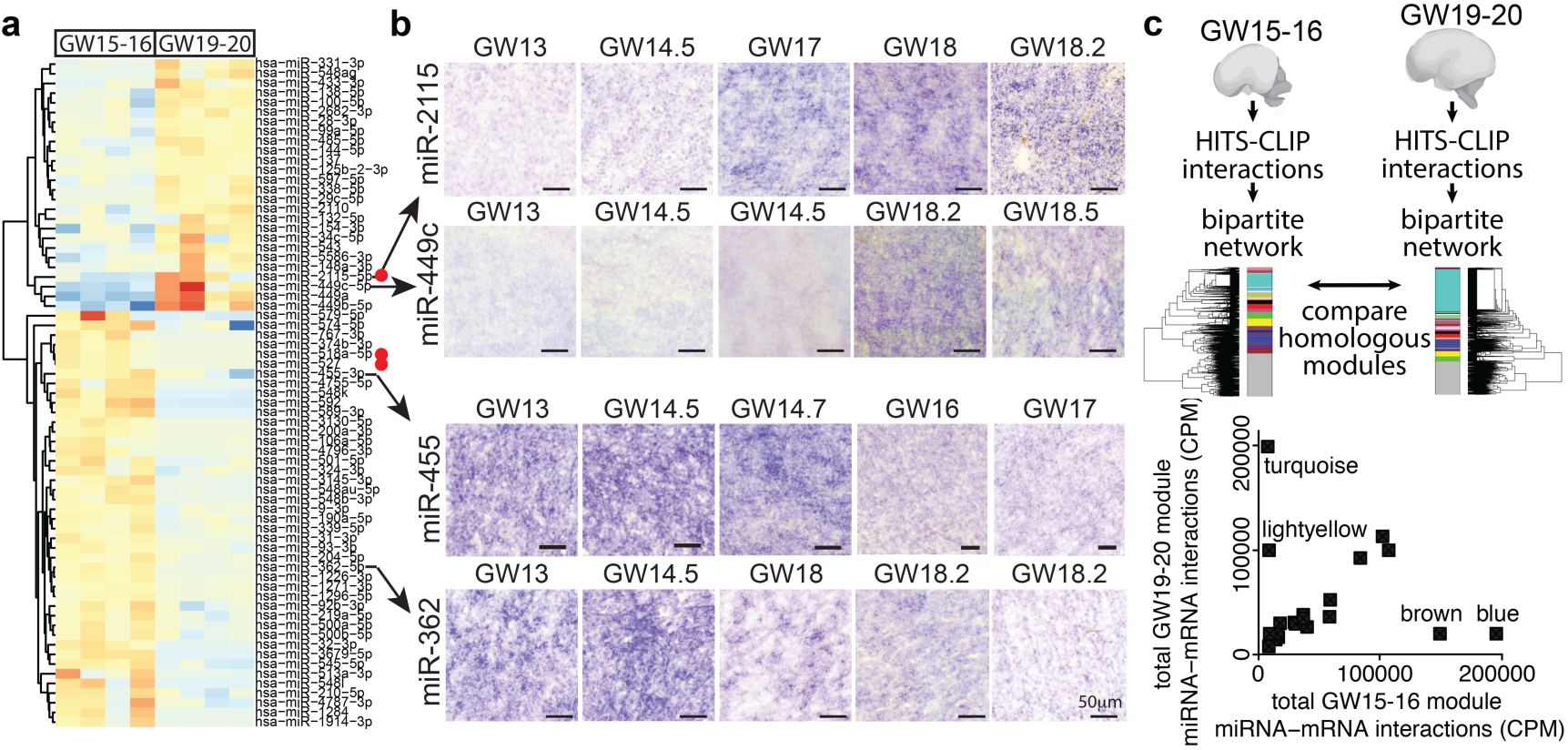
Dynamic changes in miRNA regulatory networks during development. **(a)** Comparison of miRNA abundance changes during development. **(b)** Validation of differentially expressed miRNAs by *in-situ* hybridization. Scale bar 50µm. **(c)** Module preservation analysis demonstrates significant similarity between most modules obtained for networks generated across GW15-16 and across GW19-20 samples. GW15-16 module names are used in the plot.

The developmental transition between GW15-16.5 and GW19-20.5 coincides with a depletion of proliferative capacity in the human ventricular zone^20^. Among the top five miRNAs differentially expressed between these stages, a great ape specific miRNA, miR-2115 shows prominent up-regulation at GW19-20 in the germinal zones (Fig. 3a-b Supplementary Fig. 8, 10) and could influence cortical progenitor cell function. Among miR-2115 targets, *ORC4*, a known regulator of DNA replication, is enriched in radial glia at early stages of development^10,13^, is a member of the turquoise module, and is enriched for GW19-20.5 interactions at miR-2115 response element (Fig. 3c, Supplementary Table 11). We hypothesized that miR-2115 acts through a radial glia enriched gene regulatory network involving *ORC4* to regulate cell cycle dynamics (Fig. 4a-b). After overexpressing a synthetic mmu-miR-2115 (see Methods) in developing mouse cortex, we found an increased proportion of radial glia, but fewer radial glia in mitosis, among the electroporated cells (Fig. 4c-e). Similarly, upon overexpression of miR-2115 in primary human cortical progenitors, cumulative BrdU incorporation revealed a significant increase in the total cell cycle length (Fig. 4f-h). These experiments suggest that miR-2115 recently evolved in great apes and integrated into post-transcriptional regulatory networks controlling cell cycle dynamics during human cortical development.

**Fig. 4.**
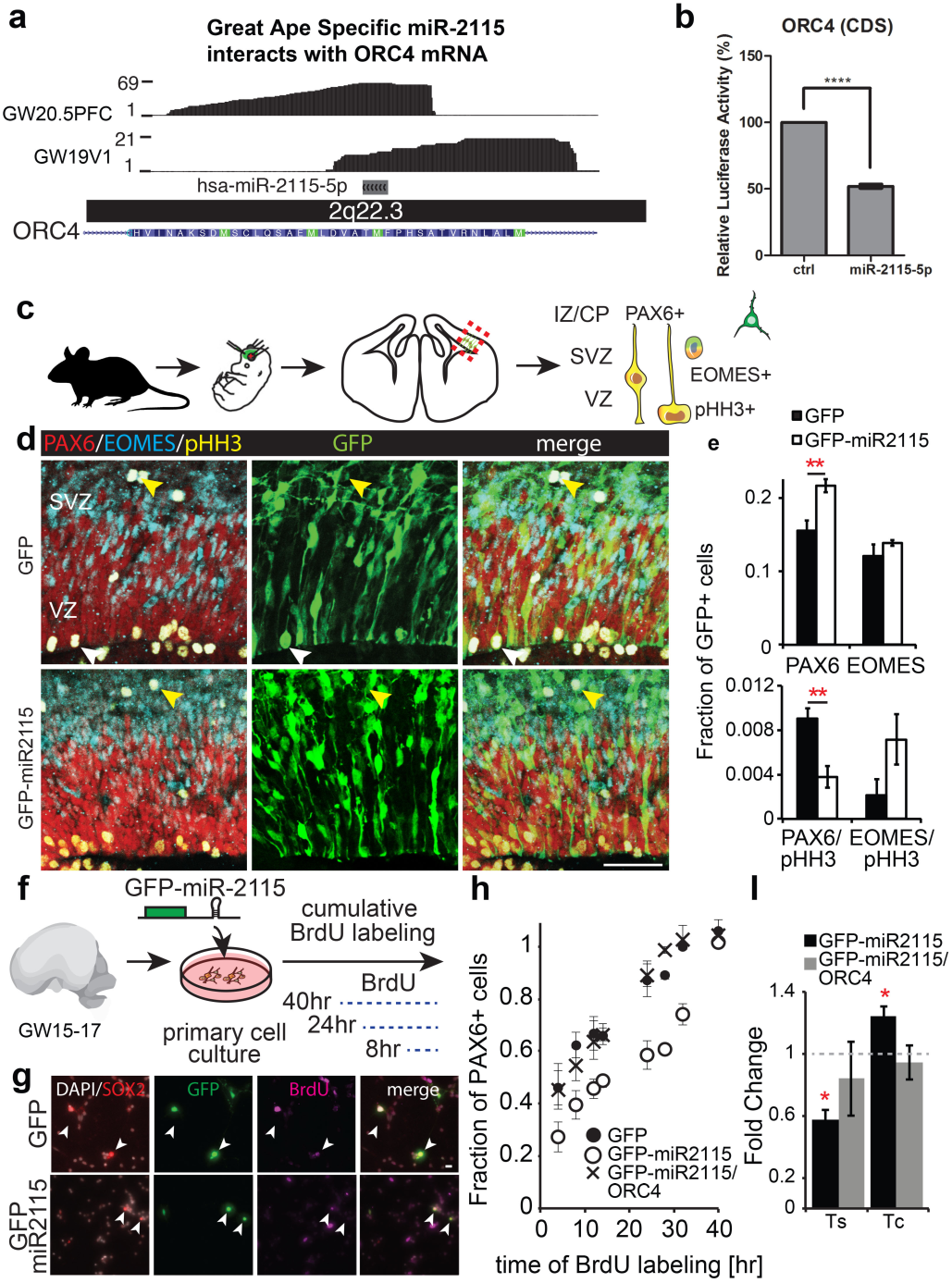
miRNAs contribute to cell-type-specific function. **(a)** Predicted miR-2115 interaction with *ORC4* mRNA in the CDS. **(b)** Luciferase reporter assay (^****^-p<0.0001). **(c)** Schematic of mouse *in vivo* experimental design. **(d)** Mouse cortex two days after electroporation at E13.5. White arrowhead indicates example of a dividing GFP+ radial glia cell, while yellow arrowheads indicate dividing intermediate progenitor cells. **(e)** Quantification of immunopositive cells. **(f)** Experimental design of cumulative BrdU labeling in primary human cells *in vitro*. **(g)** Immunostaining of human cultured primary cells. Arrowheads indicate GFP and SOX2 double positive cells. **(h-i)** Quantification of BrdU labeling of PAX6 positive cells **(h)** and estimates of S-phase length (Ts) and cell cycle length (Tc) **(i)** relative to control conditions (N = 3 biological replicates).

Our study reveals three distinct mechanisms by which miRNA regulatory pathways contribute to human brain development. Our novel single cell profiling approach revealed dynamic regulation and cell type specificity of co-expressed miRNA modules during neuronal differentiation, suggesting that miRNAs promote cell fate transitions in the cortex. In addition, by intersecting high throughput profiling of miRNA-mRNA interactions with cell type specific gene expression profiles, we can validate cell type specific miRNA/mRNA interactions and consequently cell type specific networks. Finally, we find that the abundance of individual mature miRNAs represents a component of cell identity not commonly captured by scRNA-seq approaches. Together, comprehensive understanding of cell-type-specific miRNA-mRNA interactions may reveal previously unappreciated patterns of selective vulnerability of cell-types in neurodevelopmental disorders, including Autism Spectrum Disorders.

## Author Contribution

KSK, NR, TJN, HRZ, LP, ARK designed and supervised the study. JAW, AL, BA designed and optimized single cell miRNA and mRNA PCR protocol. BA, NR, TJN, AAP, BA performed experiments. HRZ, MG, KH, BA, NR, TJN performed data analysis. TJN, NR, KK wrote the paper with contribution from all authors.

## Acknowledgement

The authors thank Shaohui Wang, Carmen Sandoval-Espinosa, Elmer Guzman, Aparna Bhaduri, Nianzhen Li for providing research resources, technical help, and helpful comments during manuscript preparation.

## Data Availability

The data used in this study are available as part of the publicly available Gene Expression Omnibus database under the accession number GSE107468.

